# The network properties of the brain at the time of normal birth support the acquisition of language processing

**DOI:** 10.1101/282673

**Authors:** Piergiorgio Salvan, Tomoki Arichi, Diego Vidaurre, J Donald Tournier, Shona Falconer, Andrew Chew, Serena J Counsell, A David Edwards

**Affiliations:** Centre for the Developing Brain, Division of Imaging Sciences & Biomedical Engineering, King’s College London, London, UK; Wellcome Centre for Integrative Neuroimaging, Oxford Centre for Functional MRI of the Brain, Nuffield Department of Clinical Neurosciences, University of Oxford, Oxford, UK; Department of Bioengineering, Imperial College London, London, UK; Wellcome Centre for Integrative Neuroimaging, Oxford Centre for Human Brain Activity, Department of Psychiatry, University of Oxford, Oxford, UK

## Abstract

Language acquisition appears to rely at least in part on recruiting pre-existing brain structures. We hypothesized that the neural substrate for language can be characterized by distinct, non-trivial network properties of the brain, that modulate language acquisition early in development. We tested whether these brain network properties present at the normal age of birth predicted later language abilities, and whether these were robust against perturbation by studying infants exposed to the extreme environmental stress of preterm birth.

We found that brain network controllability and integration predicted respectively phonological, ‘bottom-up’ and syntactical, ‘top-down’ language skills at 20 months, and that syntactical but not phonological functions were modulated by premature extrauterine life. These data show that the neural substrate for language acquisition is a network property present at term corrected age. These distinct developmental trajectories may be relevant to the emergence of social interaction after birth.

## Main text

Language acquisition appears to rely at least in part on the recruitment of pre-existing brain structures and capacities ^1–3^: during the last trimester of pregnancy temporal brain areas are active during passive language listening ^4–6^; ventral and dorsal white-matter bundles are present during early infancy ^7,8^; and over the first months of postnatal life, infants progress from a general capacity to discriminate any phonetic contrast to a more specific response to the combination of vowels and consonants of their native language ^9–11^. The establishment of a phonetic repertoire allows infants progressively to infer statistical patterns in speech and language structure ^12,13^ and later progress to the first production of words at 9-12 months of age and multiword utterances and grammar by 18-24 months of age^2^. Before children start to read, the pre-existing connectivity profile of the visual word form area predicts its future location ^3^, thus also suggesting that cognitive brain development also takes advantage of pre-existing cortical circuits ^14^.

However, the evolution of a neural substrate for language during the perinatal period remains incompletely understood. Brain development is extremely rapid at this time and normal maturation is significantly disrupted by environmental pressures such as preterm birth which lead to widespread abnormalities in structure and connectivity, including substantial changes within regions involved in language processing ^15,16^, which significantly predict later language function. Nevertheless despite these significant abnormalities, the effect of early adversity on language function is complex. Language acquisition is frequently disrupted by preterm birth. However the effect is complex in preterm infants: as whilst earlier exposure to speech does not alter language perception in early childhood ^17^, they show increasing risk of impairments later in life ^18–21^. This suggests that some aspects of language acquisition are more susceptible to environmental pressures during the preterm period than others.

From 30 to 44 weeks gestational age brain network architecture matures rapidly, and at term corrected age the brain shows scale-free macroscopic structural connectivity largely similar to the adult ^22,23^. Although widespread abnormalities are induced by preterm birth, this is less evident on the connections between the hub nodes ^22^ rather than the feeder nodes of the network ^22,24^, which therefore predicts robust network functionality is retained despite widespread damage. There is also increasing evidence that multisensory input and presumably processing across different brain systems are integral to language acquisition in the perinatal period ^25^. This combination of factors led us to hypothesize that in preterm infants without focal destructive brain lesions, the neural substrate for language is in part a network property of the network.

Here, we examine language development in the perinatal period as properties of the large-scale structural connectivity network ^26^, testing the prediction that network controllability and integration are critical features in effective language acquisition early in development. Studying a group of infants born preterm allowed us to investigate the effect of adverse environmental stimuli on these language network properties. We found that language abilities in early childhood are predicted by network metrics at the normal time of birth. We show that ‘bottom-up’ phonological processing is unaffected by preterm delivery while ‘top-down’ syntactic processing is altered by premature extra-uterine exposure. These results show that the neural substrate for language is at least in part a network property present at term corrected age and suggest a mechanism whereby language acquisition may support the development of social behaviors.

## Results

We acquired high b value, high angular resolution diffusion weighted-magnetic resonance imaging (DW-MRI) from 43 preterm born infants (median gestational age (GA) at birth of 30 weeks) at term equivalent age (median postmenstrual age (PMA) at scan of 42 weeks) (Table S1). At 20 months of age, we formally assessed their level of cognitive and linguistic abilities. The whole-brain white-matter connectome was reconstructed using anatomically constrained probabilistic tractography ^27^, based on a 90-region parcellation from the Infant Brain Atlas for Neonates ^28^ (*see Materials and Methods*) (Fig. 1).

**Figure 1:**
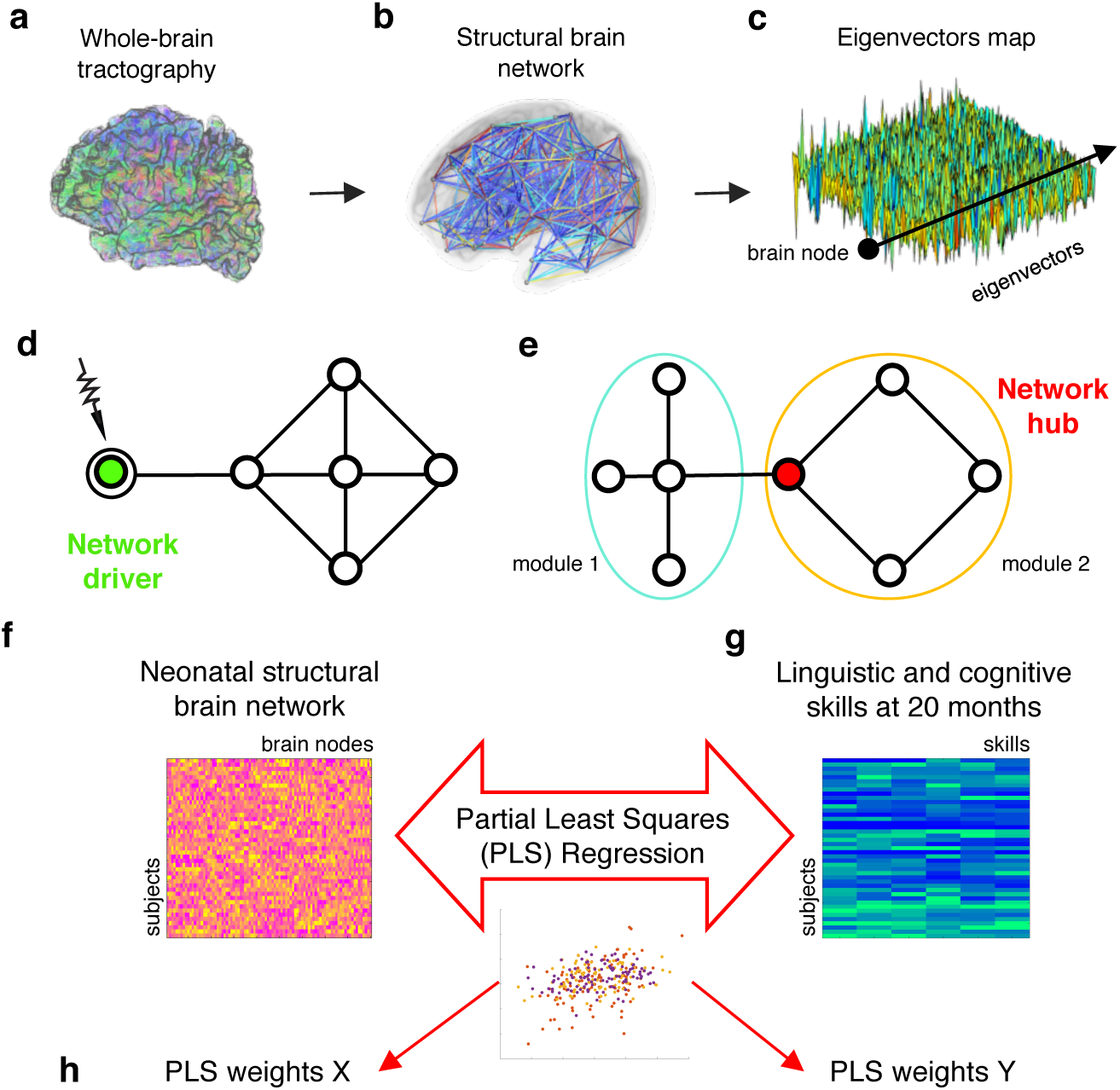
Exploring the link between neonatal brain network properties and the acquisition of language in childhood. 43 preterm born infants were imaged at the time of normal birth using high *b* value, high-angular resolution diffusion-weighted MRI. **(a)** Whole-brain streamline tractography was performed, and the **(b)** neonatal brain structural network was reconstructed to study its emerging properties. **(c)** The involvement of each network node across the network eigenvectors was characterized. This was then used to study: **(d)** *Structural controllability*, which predicts individual nodes (or *drivers*) that are theoretically able to change a network’s activity state. In this example network, the node highlighted in green is a *network driver* which can change network nodes activity and steer the system to a different state. **(e)** *Network integration*, characterizes the diversity of intermodular connections, and thus identifies individual nodes (or *hubs*) that are likely to facilitate global intermodular integration. In the example network, the node highlighted in red is a *network hub* which integrates information between two distinct network modules (circled in blue and orange). We then used Partial Least Square regression to associate neonatal brain network properties **(f)** with linguistic and cognitive skills assessed at 20 months of age **(g)**. Significant models, as well as significant model weights, were then characterized in order to quantify the brain network features related to language acquisition.

We used commonly used network metrics ^26^ and cross-validated multivariate models to identify the relationship between the computational properties of connections within the neonatal brain with linguistic and cognitive abilities in later childhood. We present significant results for those brain-language models after Bonferroni correction for multiple comparison. A comprehensive quantitative comparison with alternative models and random brain networks is reported in the *Supplementary information* (*SI*).

### Neonatal brain network controllability predicts phonologic language abilities in childhood

Diffusion MR imaging provides a rich and reproducible mapping of the macroscopic white matter connectivity network in the brain. However this mapping is not itself sufficient to understand the role of the network in neurological functions, for which further analysis of the network properties are needed ^29^. To understand whether language emerges from pre-existing connectivity patterns, we therefore asked if inter-subject differences in the perinatal structural brain network properties relate to the development of language skills in childhood?

To study the relationship between the neonatal brain’s structural connectome and how its topology facilitates transition from one activity state to another, we analyzed network controllability ^30–32^. Given constraints imposed by the brain’s underlying structural connections, this approach allowed the identification of network “controller” or “driver” sets, comprising network nodes (in this case specific brain regions) which are critical to guide the system to the activity states required for the acquisition of later language abilities. This then allowed testing of the hypothesis that synergic activity in sets of separate brain regions ^33^ can explain differences in language behavior.

We derived a simplified linear model from the underlying structural brain network ^34,35^ (*see Materials and Methods*) and quantified structural *modal* controllability (here referred to as structural controllability) for each brain region, predicting the ability of a node to guide the system’s entire dynamics ^36,37^. Consistent with previous work which suggests that driver nodes are usually not high-degree “hub” regions, we found that structural controllability was inversely related to weighted degree (or nodal strength, the sum of the weights of the edges; Pearson’s rho = −0.94) (Fig. S1) ^32^. We also found that subcortical structures and the orbitofrontal cortex have the highest structural controllability scores.

To identify sets of controllers linked with the development of language skills we applied cross-validated partial least-squares (PLS) regression ^38,39^ to identify linear combinations of the independent variables (in this case: brain network metrics) that maximally predict linear combinations of the dependent variables (in this case: behavioral language measures).

Having regressed out the effect of possible confounding clinical and sociodemographic factors from both the imaging data (PMA at scan; degree of prematurity (GA at birth); and sex) and behavioral measures (socioeconomic score (SES), GA at birth and sex) we found significant modes of covariation between patterns of nodal controllability at the neonatal time-point with measures of language and cognitive abilities at 2 years of age. We found that inter-individual differences in structural controllability were significantly associated with inter-individual differences in a latent factor of language abilities, (Adjusted-R^2^ = 0.64; p-value = 5 × 10^−4^; Mean Squared Error (MSE) = 1.9282; p-value = 5 × 10^−4^; predicted-R^2^ = 0.27; p-value = 1.6 × 10^−3^; Mean Squared Predicted Error (MSPE) = 5.5528; p-value = 1.6 × 10^−3^; all significance values were compared against 10,000 permutations) (Fig. 2, Table 1, 2, Fig. S2). This relationship shows that subjects who exhibited higher structural control in the left superior and middle-temporal cortex were those who later exhibited greater vocabulary scores and more advanced words combination capacities. This combination of particular language abilities and brain regions, suggest that this latent factor may rely predominately on phonological skills or more general bottom-up processing of auditory-verbal material ^40^. This finding suggests that in the neonatal brain the left temporal and supramarginal cortices represent a network controller set which support the dynamic control of the functional brain activity which underlies later language development.

**Figure 2:**
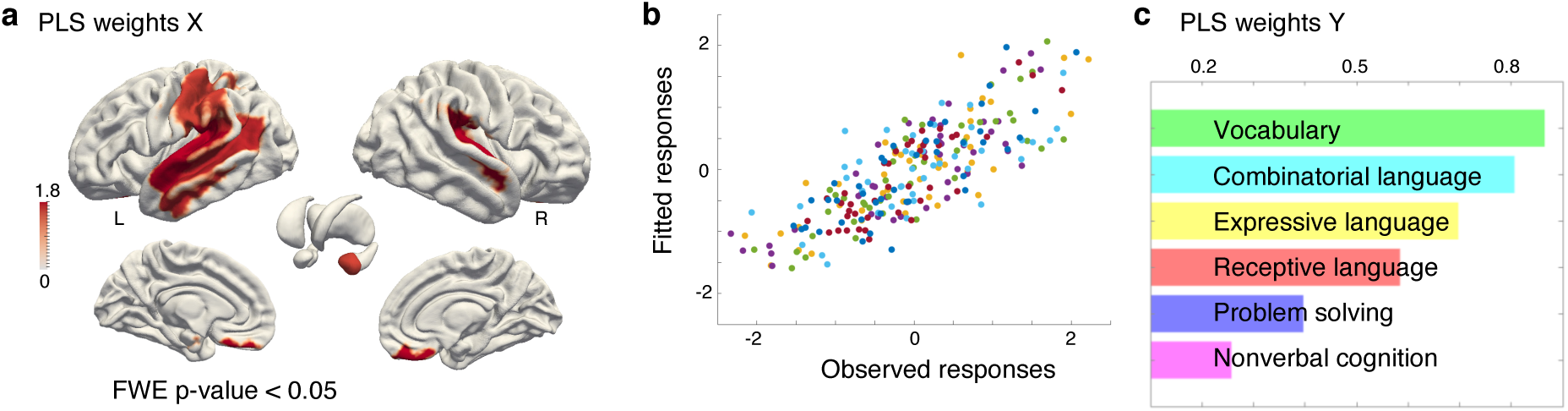
The left superior and middle-temporal cortex is network driver underlying the development of bottom-up language processing. We found that structural controllability in the neonatal brain predicts bottom-up language abilities in childhood. In the identified significant relationship, neonates who exhibited significantly greater structural controllability scores at the neonatal timepoint in the left superior and middle-temporal cortex (PLS weights X, **(a)**), were those who later developed a larger vocabulary and more advanced language combinatorial capacities (PLS weights Y, **(c)**). The association between the observed and fitted behavioral skills **(b)** was highly significant (Adjusted-R^2^ = 0.64; p-value = 5 × 10^−4^; colors in barplot **(c)** are matched with behavioral skills in scatterplot **(b))**. All of the significant brain network regions shown in **(a)** are also summarized in (Table 2a). Together these results suggest the left temporal lobe acts as a network driver of the brain activity underlying the acquisition of phonological abilities.

**Table 1:**
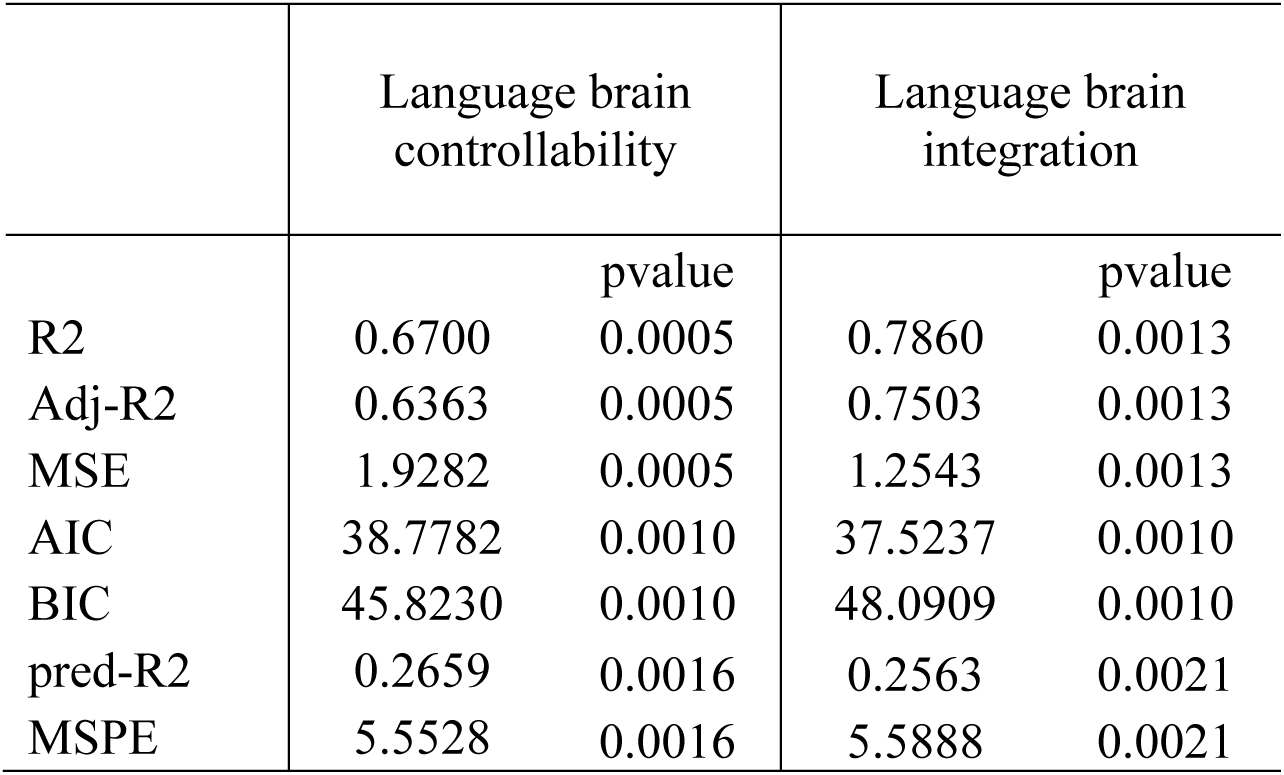
Fit and prediction results in neonatal brain models of language acquisition. Shown are the results of Partial Least Square (PLS) fitting and prediction for language brain controllability and language brain integration. PLS regression models were based on 10-folds cross-validation with 100 Monte-Carlo repetition and 10,000 permutations of response variable. Models were assessed on the basis of: adjusted R^2^ (Adj-R2); Mean Squared Error (MSE); Akaike information criterion (AIC), and Bayesian information criterion (BIC), predicted R2 (pred-R2); cross-validated Mean Squared Predicted Error (MSPE).

|  | Language brain controllability |  | Language brain integration |  |
| --- | --- | --- | --- | --- |
|  |  | pvalue |  | pvalue |
| R2 | 0.6700 | 0.0005 | 0.7860 | 0.0013 |
| Adj-R2 | 0.6363 | 0.0005 | 0.7503 | 0.0013 |
| MSE | 1.9282 | 0.0005 | 1.2543 | 0.0013 |
| AIC | 38.7782 | 0.0010 | 37.5237 | 0.0010 |
| BIC | 45.8230 | 0.0010 | 48.0909 | 0.0010 |
| pred-R2 | 0.2659 | 0.0016 | 0.2563 | 0.0021 |
| MSPE | 5.5528 | 0.0016 | 5.5888 | 0.0021 |

**Table 2:** Brain network regions significantly involved in supporting acquisition of language processing. Summary of brain network regions with significant PLS weights (FWE-corrected p-value < 0.05 across all brain regions), respectively for language brain controllability **(a)** and language brain integration **(b)**. These regions are also shown in Figure 2 and 3.

| Region | AAL Abbreviation | PLS weights X (W*) | FWE pvalue |
| --- | --- | --- | --- |
| Orbitofrontal cortex (medial) right | ORBmed-R | 1.8667 | 0.0235 |
| Amygdala left | AMYG-L | 1.5744 | 0.0472 |
| Inferior parietal lobule left | IPL-L | 1.5816 | 0.0467 |
| Supramarginal gyrus left | SMG-L | 1.5711 | 0.0473 |
| Superior temporal gyrus left | STG-L | 1.9501 | 0.0194 |
| Superior temporal gyrus right | STG-R | 1.8982 | 0.0212 |
| Middle temporal gyrus left | MTG-L | 1.701 | 0.0343 |

(b)
| Region | AAL Abbreviation | PLS weights X (W*) | FWE pvalue |
| --- | --- | --- | --- |
| Middle frontal gyrus left | MFG-L | 2.5161 | 0.0040 |
| Inferior frontal gyrus (triangular) left | IFGtriang-L | 1.6908 | 0.0360 |
| Middle occipital gyrus right | MOG-R | 2.6982 | 0.0025 |
| Temporal pole (middle) left | TPOmid-L | 2.2326 | 0.0089 |
| Temporal pole (middle) right | TPOmid-R | 1.9949 | 0.0175 |

### Neonatal brain network integration predicts top-down language abilities in childhood

We then studied network brain integration to assess how the connectivity profile of an individual brain region can facilitate communication between groups of densely interconnected nodes (modules) across the brain. This is thought to represent the ability of a single brain area to rapidly combine specialized information from distributed regions ^41^ and may represent a mechanistic property through which the brain creates complex, high-order, adaptive behavior ^42,43^.

In adults, brain cognitive systems are hierarchically distinct based on their level of network integration ^26^. Here, we calculated network integration as the *participation coefficient* (*see Materials and Methods*), measuring the diversity of inter-modular connections of individual nodes ^41,42,44^ (Fig. S1). Regions with greater values of participation coefficient (close to one) have connections uniformly distributed among all of the modules, whilst regions with smaller values (towards zero) have all of their connections within their own module. We hypothesized that network integration is an important property for neurocognitive development, and asked how network integration in the neonatal brain represents an early neural basis for the development of language skills in early childhood.

In the neonatal brain, we found that the network drivers of activity are not necessarily also brain network connector systems, as we found that network integration was not related to structural controllability (Pearson’s r = −0.01).

By applying the same cross-validated PLS regression approach described above, we found that higher language performance in childhood was predicted by greater network integration in the left inferior, medio-prefrontal cortex, the temporal poles and right occipital gyrus (Adjusted-R^2^ = 0.75; p-value = 1.3 × 10^−3^; MSE = 1.2543; p-value = 1.3 × 10^−3^; predicted-R^2^ = 0.26; p-value = 2.1 × 10^−3^; MSPE = 5.5888; p-value = 2.1 × 10^−3^) (Fig. 3, Table 1,2, Fig. S3). Subjects who exhibited higher network integration in the left prefrontal cortex were those that later demonstrated greater understanding of pronouns and sentences in the past tense; and later showed greater naming abilities and greater production of present progressive form or multiple-word questions. These results suggest that the prefrontal cortex acts as a network connector system which is essential for the development of later syntax and semantic language skills ^1,2,11^, in accordance with adult studies which have found that greater integration across specific networks are related to higher cognitive performance ^45^.

**Figure 3:**
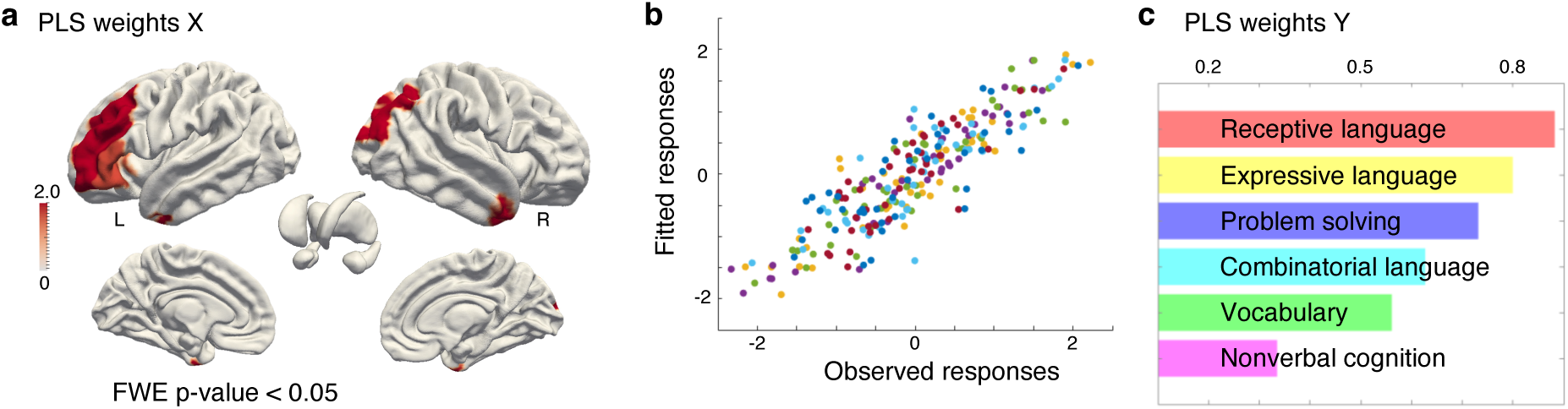
The left inferior and middle-prefrontal cortex is a network hub which supports the development of top-down language processing. We found that network integration in the neonatal brain predicts top-down linguistic abilities in childhood. In this relationship, neonates who had significantly greater network integration scores at the time of normal birth in the left inferior and medio-prefrontal cortex (PLS weights X**(a)**), were those who developed more advanced receptive and expressive language abilities in semantics and syntax (PLS weights Y **(c)**). The association between the observed and fitted behavioral skills was highly significant (Adjusted-R^2^ = 0.75; p-value = 1.3 × 10^−3^; colors in barplot **(c)** are matched with behavioral skills in scatterplot **(b)**). All of the significant brain network regions shown in **(a)** are also summarized in (Table 2b). These results provide the first quantitative evidence that the left prefrontal cortex at the time of normal birth has organized structural connectivity which aids the later development of top-down linguistic abilities.

### Impact of early environmental exposure on brain mechanisms of language acquisition

To answer the question of whether *early extra-uterine exposure* affects the identified neonatal brain network mechanisms for language acquisition, we tested whether the *degree of prematurity* modulates the proposed models of language brain controllability and language brain integration. We estimated brain-behavior PLS regression models without co-varying for the effect of GA at birth, and then split the sample into two groups of preterm infants with low and high GA at birth (Fig. S4). We then tested two separate classifiers based on the PLS individual subject weights which represent the relationship between the brain and the behavior measures (*see Materials and Methods*) and tested the effect on network controllability and integration measures separately. Using cross-validated linear discriminant analysis (LDA) we found that *only* when using the PLS subject weights derived from language brain integration could the separation between preterm infants with low and high GA be significantly predicted (respectively: language brain integration, accuracy = 0.74; p-value = 4.6 × 10^−3^; language brain controllability, accuracy = 0.58; p-value = 0.33; significance of classification was assessed against 10,000 permutations of individual subject weights) (Table 3).

**Table 3:** Language brain integration discriminates between infants with high and low gestational age at birth. To understand whether the degree of prematurity has an effect on language brain controllability and/or integration, we trained and tested PLS individual subject weights (or PLS score pairs: XS, YS) to discriminate between neonates with low vs high GA at birth. We found that language brain integration (but not language brain controllability) carries information which significantly discriminates between infants with GA at birth lower or greater than 29 weeks, suggesting an effect of degree of prematurity specifically in this brain mechanisms for language acquisition.

|  | XS,YS Modal controllability (not adjusted for GA) | XS,YS Network integration (not adjusted for GA) |
| --- | --- | --- |
| Accuracy | 0.5814 | 0.7442 |
| pvalue | 0.3260 | 0.0046 |

We then explored how language brain integration was different between infants with low and high GA at birth by estimating a PLS model relating network integration and behavior measures separately for each group. Using a T-test, we assessed group-differences in the *PLS coefficients* that relate the original sets of brain and behavior variables to the identified PLS latent components (formally the PLS weights W*; *see Materials and Methods*). We found that the only significant differences were in the right superior frontal gyrus and in the right rolandic operculum (Family-wise error (FWE) corrected p-value, respectively = 2.5 × 10^−2^ and 3.6 × 10^−2^). This result shows that preterm infants with low GA at birth have greater language network integration in the right hemisphere, suggesting those delivered earlier have a more bilateral pattern of language brain architecture at term equivalent age.

We further tested whether the modulation of degree of prematurity in the model of language brain integration could be simply explained by the correlation between GA at birth and the *brain measures* (regardless of behavior). Using a *univariate* general linear model (GLM), we found that there was not a significant relationship between GA at birth and the regional measures of network integration (either one-tail or two-tails T-test; FWE-corrected p-values > 0.7005, tested against 10,000 permutations). We also found that *behavioral measures* could not explain the modulation language brain integration by GA at birth (univariate GLM testing; FWE-corrected p-values > 0.1730). Finally, we also tested whether *multivariate* patterns of brain or behavior measures could significantly discriminate between preterm infants with low and high GA at birth. Using the LDA, we also found no evidence for this relationship (brain measures: accuracy = 0.58; p-value = 0.30; behavioral measures: accuracy = 0.56; p-value = 0.40; significance of classification was assessed against 10,000 permutations brain or behavior measures).

These additional analyses suggest that the modulation of language brain integration by the degree of prematurity is not due to the effect of premature birth on a single brain region or cognitive skill. Instead, premature birth appears to affect the complex brain network which underlies higher-order language abilities.

## Discussion

These results demonstrate that distinct childhood language abilities can be predicted by specific network metrics in the perinatal period. This suggests that the brain’s structural network is intrinsically organized to predispose itself for language acquisition. We further found that premature environmental exposure significantly affected the relationship between network features and syntactic but not phonological abilities, suggesting distinct neurodevelopment trajectories for top-down and bottom-up human language acquisition.

### A structural brain network underpins efficient language learning

We show that even at the time of normal human birth, the brain’s structural network already possesses specific patterns of controllability and integration within regions known to play a fundamental role in language function throughout the lifespan.

We first studied structural network controllability which predicts the ability of individual nodes (or *drivers*) to theoretically steer the network’s activity output. We found that the left temporal and supramarginal cortices were key drivers of activity across the network which predict inter-subject differences in childhood language skills. In keeping with this finding, the temporal regions have been shown to be activated in an adult-like fashion during passive listening of phoneme deviants and human speech even before and around the time of normal birth ^5,6^. Here, we additionally show that language brain network controllability represents an early form of structural specialization which supports the bottom-up, phonological learning of auditory-verbal material, and may represent an early substrate for auditory-verbal working memory ^40^.

We then studied network integration, to measure the ability of individual nodes (or *hubs*) to facilitate global intermodular connectivity. In this context, we found that the left prefrontal cortex, the temporal poles, and the right occipital cortex, are key parts of a brain hub system responsible for the development of receptive and expressive linguistic skills and problem solving. Previous studies have shown that the left prefrontal region responds to speech presentation by three months of age ^4^, and that some degree of lateralization in response to phoneme deviance is already present in this area in the preterm brain even before term equivalent age ^6^. Although it was previously thought that the left inferior frontal gyrus does not show selectivity for syntax until 9-10 years of age ^1,46^, our findings provide evidence that structural connectivity for top-down, syntactic and semantic abilities is established as a network property as early as at the normal time of birth, preceding the emergence of functional specialization and its behavioral correlate. It is also possible that these network properties may aid the learning of reading skills, through mediating the neuroarchitectural integration of the vision and language processing areas ^14^.

Together, these findings demonstrate the presence of a structural brain network that is already organized to support environmental interaction and language learning. Of particular significance, these results are in agreement with brain network models of mature language processing and development. The left inferior and middle-frontal regions, and the superior and middle-temporal cortices are known to alter their activity during auditory-verbal working memory tasks ^47^ and are hypothesized to represent the brain basis underlying the learning of new words and unfamiliar phoneme combinations ^40^. This left language network is also thought to support core syntactic language processing in sentence-level argument structure^48,49^. The left inferior frontal gyrus has also been postulated to specifically modulate top-down processing of sentence-level syntax ^50,51^ by regulating the conversion of lexical and phonological information received from the superior temporal cortex via the arcuate fasciculus into an articulation code ^48,52,53^.

Structural controllability in this context may reflect a network property that is complementary to network integration. It is possible that direct long-distance connections already present at this age ^8^ enable higher-order prefrontal areas to perform top-down modulation on structural controllers such as the temporal regions, in order to ensure efficient language learning. The presence of these two network properties at the time of normal birth may represent a required computational pathway before specific refinement through experience dependent plasticity during critical windows for language acquisition in early childhood ^54^.

### Premature environmental exposure alters brain networks that are key for syntactic language acquisition

Although the posterior temporal areas have already started to develop the neural organization that underlies language processing before term equivalent age ^6^, earlier exposure to speech in preterm infants does not lead to accelerated development of phonological skills ^17^. Our results expand on this further by confirming that the neonatal brain network features which predict bottom-up language abilities in later childhood are minimally influenced by early environmental exposure. In contrast, we found that the prefrontal brain network architecture which predicts childhood top-down language skills such as semantic and syntactic language learning are significantly altered by premature environmental exposure. This may be because such processes are computationally more expensive than phonology and rely on connections that are not as well developed in the perinatal period as those of the temporal cortices. These findings are consistent with those of our previous study which showed that preterm birth affects the structural connectivity of sparsely-connected areas, whilst minimally impacting those related to hub regions ^22^.

Preterm-born children are known to have difficulties in complex language attainment and function which increase with age ^55,56^. These may result from general cognitive difficulties rather than a specific phonological impairment ^20,57^, as higher-order semantic and syntactic knowledge is thought to involve hierarchical integration across language areas and a significant working memory component ^1^. The results of this study show that premature environmental exposure alters the prefrontal neonatal brain network structure underlying future, top-down language learning.

Although structural network controllability represents an important step towards a novel mechanistic understanding of the human brain’s structure-function relationship ^26,29^, it is nevertheless important to acknowledge the limitations of this framework ^36,37^. Although system brain dynamics may be non-linear in nature to some degree, the framework we used relied on linear modelling as such methods have been successfully applied previously to predict dynamics in the adult human brain ^35^. The assumption of linear dynamics is nevertheless true when close to operating points and over short time horizons ^36,37^.

### Conclusions

In this work, we provide evidence that at the normal time of birth, in addition to known local and areal mechanisms, the neural basis for language acquisition includes properties of the overall structural brain network, and that network controllability and integration respectively underlie bottom-up and top-down language skills in childhood. We further show that preterm birth leads to specific impairment these properties that affect future semantic and syntactic learning. This work provides insight into the fundamental relationship between the early establishment of human brain network architecture and complex behavior, and has implications for fostering healthy long-term outcomes and social interaction.

## Materials and methods

### Infants

Preterm infants were recruited as part of the Evaluation of Preterm Imaging study (Eprime), and were imaged at term equivalent age over a 3 year period (2010-2013). The study was reviewed and approved by the National Research Ethics Service, and all infants were studied following written parental consent. A cohort of 43 preterm born infants (median age at birth of 30.14 gestational (GA) weeks; range 24 - 32; 18 females) with no evidence of focal abnormality on MRI were imaged using high-angular resolution diffusion-weighted neuroimaging at 42.14 postmenstrual (PMA) weeks (range 39 - 46).

Standardized neurodevelopmental assessment at a median age of 22 months (range: 21 – 24 months; median of 20 months corrected for prematurity) was carried out by experienced pediatricians with the *Bayley III scales of infant and toddler development* (BSID-III) ^58^ and of the *Parent Report of Children’s Abilities Revised* (PARCAR) ^59^. None of these behavioral scales were significantly correlated with GA at birth or PMA at time of MRI scan. As previously described in the literature, there was a significant effect of socioeconomic score ^60^. The BSID-II measures the actual cognitive and linguistic abilities that the child is able to demonstrate to the examiner during the testing session. Specifically, the receptive communication subscale aims to measure receptive language development, with items covering vocabulary and semantics (i.e. picture and object series identification; understanding of inhibitory words), and morphology and syntax (i.e. object categories identification; understanding of plurals; understanding of pronouns; understanding of past tense). Whilst the expressive communication subscale aims to test expressive language development, with its items measuring semantics (i.e. the use of one-word approximations; naming action picture series), morphology, and syntax (i.e. production of present progressive form; production of prepositions; explain how an object is used; production of multiple-word questions). The cognitive scale aims to measure i.e. attention, problem solving, reasoning, memory, object novelty, two-step actions, size discrimination, number sequences repetition, and representational play. The PARCAR measures the level of behavioral capacities reported by the parents or guardians. 34 items are summed to give a nonverbal cognitive subscale score. Whilst linguistic development is assessed with 2 subscales that are part of the MacArthur Communicative Development Index Words and Sentences ^61^: a 100-word vocabulary checklist and 18 items to assess grammatical competence.

### Acquisition of MRI imaging data

All MRI studies were supervised by an experienced pediatrician or nurse trained in neonatal resuscitation. Pulse oximetry, temperature, and heart rate were monitored throughout the period of image acquisition; hearing protection in the form of silicone-based putty placed in the external ear (President Putty, Coltene; Whaledent) and Mini-muffs (Natus Medical Inc.) was used for each infant. Sedation (25–50 mg/kg oral chloral hydrate) was administered to 33 infants. Imaging was acquired using an eight-channel phased array head coil on a 3-Tesla Philips Achieva MRI Scanner (Best, Netherlands) located on the Neonatal Intensive Care Unit. Whole-brain diffusion-weighted MRI data were acquired in 64 non-collinear directions with b value of 2500 s/mm^2^ and 4 images without diffusion weighting (isotropic voxel size of 2 mm; TE= 62 ms; TR= 9000 ms). High-resolution anatomical images were acquired with pulse sequence parameters: T1 weighted 3D MPRAGE: repetition time (TR) = 17 ms, echo time (TE) = 4.6 ms, flip angle 13°, slice thickness 0.8 mm, field-of-view 210 mm, matrix 256 × 256 (voxel size: 0.82 × 0.82 × 0.8 mm); and T2 weighted fast-spin echo: TR = 8670 ms, TE = 160 ms, flip angle 90°, slice thickness 2 mm with 1 mm overlap, field-of-view 220 mm, matrix 256 × 256 (effective voxel size: 0.86 × 0.86 × 1 mm).

### MRI data pre-processing

Structural MRI volumes were processed in order to extract tissues segmentation ^62^. Diffusion MRI volumes were visually inspected and processed. B0 field inhomogeneities, eddy currents, and inter-volume motion were corrected and outlier slices were replaced using topup and eddy tools in FSL5 ^63–65^. B1 field bias was corrected using ITK-N4 ^66^. All rigid registrations in native subject space were estimated using FSL boundary-based registration optimized for neonatal tissue contrasts ^67–69^; and nonlinear registrations to the T2-weighted template were estimated using Advanced Normalization Tools (ANTs) ^70^. All transformation pairs were calculated independently and combined into a single transform in order to reduce interpolation error. This preprocessing approach was previously published ^8^.

### Neonatal connectome

Estimation of fiber orientation distribution was computed through constrained spherical deconvolution with maximum spherical harmonic order of 8 ^71^. Whole-brain fiber tracking was performed using *anatomically constrained* probabilistic tractography from the MRtrix3 package with 10 million streamlines ^27,72^ and SIFT2 regularization ^73^. Subject-specific weighted adjacency matrices of the brain connectome were computed based on a 90-region neonatal atlas ^28^ and edge weights were defined by the number of streamlines connecting each pair of nodes ^36^.

### Dynamic model of neural function

Drawing from previous work predicting nonlinear brain dynamic processes using simplified linear models ^34,35^, we employed a noise-free linear discrete-time and time-invariant network model:

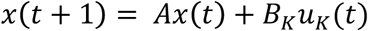

based on the underlying structural connectome, here *x*: ℝ_≥_0__ → ℝ^*N*^ describes the state of activity of brain regions over time, and *A* ∈ ℝ*^N^*^×*N*^ is the structural white-matter connectome reconstructed with diffusion neuroimaging tractography streamlines. The input matrix *B_K_* identifies the control points *K* in the brain where *K* = {*k*_1_,…, *k_m_*} and *B_K_* = [*e_k_*_1_ … *e_km_*], and *e_i_* denotes the *i^th^* canonical vector of dimension *N*. The input *u_k_*: ℝ_≥_0__ → ℝ*^m^* denotes the control strategy.

### Network controllability diagnostics

We studied the controllability of this dynamical system, the capacity of steering the state of the system to a specific target state by means of an external control input ^74^. In order words, the capacity of manipulating the dynamic trajectories of a system.

Classic results in control theory ensure that controllability of the dynamical network model previously explicated from a set of network nodes *k* is equivalent to the controllability Gramian *W_k_* being invertible, where

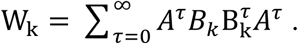

In accordance with ^36^, we applied this framework to choose controls nodes one at a time, and thus the input matrix *B* in fact reduces to a one-dimensional vector.

We studied two different quantitative, regional control strategies that describe the ability to move the network into different states.

### Structural average controllability

Structural *average* controllability describes the ease of transition to many states nearby on an energy landscape (low-energy state transitions) ^36^. In accordance with previous work ^30,75^, we define Trace(*W_k_*) as a measure of average controllability, encoding the energy of the network impulse response or, equivalently, the network H_2_ norm. This metric quantifies a node’s capacity in moving the system to many easily reachable states and equals the average input energy from a set of control nodes and over all possible neurocognitive configurations ^76^.

### Structural modal controllability

Structural *modal* controllability assesses the capacity of a node to control each evolutionary mode of a dynamical network ^77^. Modal controllability is computed from the eigenvector matrix *V* = [*v_ij_*] of the network adjacency matrix A. By extension from the Popov-Belevitch-Hautus (PBH) eigenvector test ^75^, if the entry *v_ij_* is small, then the j^−*th*^ mode is poorly controllable from the node *i*. In accordance with ^30^, we adopt

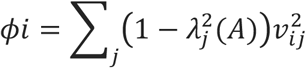

as a scaled measure of the controllability of *N* modes *λ*_0_(*A*),…, *λ*_*N*−1_(*A*) from the brain region *i*, where *λ*(*A*) are the eigenvalues of *A*. This metric quantifies a node’s capacity to control trajectories to all dynamical states of the network ^77^, hence identifying nodes sets controlling dynamic transition towards hard-to-reach (high-energy state) neurocognitive configurations.

### Multi-scale community detection

In order to quantify network integration and segregation, we first identified brain network modules: groups of densely interconnected network nodes, which often are the basis for specialized subunits of cognitive processing ^78,79^. Based on a group-level neonatal brain connectome (normalized average), we performed multi-scale community detection across a range of structural resolution parameter γ ^36,80,81^. By maximizing the modularity quality function ^82^ using a Louvain-like ^83^ locally greedy algorithm ^84^ (with 100 optimizations) for multiple values of γ, we quantitatively estimate consensus between partitions calculating partitions similarity as z-score of the Rand coefficient ^85^. Maximum mean partition similarity was achieved for γ = 4.8, hinting at the presence of 18 especially well-defined modules.

### Network integration

Network integration or between-module connectivity, was calculated as (weighted) participation coefficient ^42,44,86^. Quantifying strength of a region’s structural interactions between modules, the participation coefficient for each region was calculated as

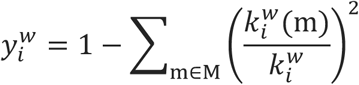

where M is the set of modules, and 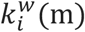 is the strengths of connections between *i* and all nodes in module *m* ^41,87^. Regions with greater values of participation coefficient (close to one) have connections uniformly distributed among all of the modules; whilst regions with smaller values (towards zero) have all of their connections within their own modules.

### Network segregation

Based on the previously identified community assignments, we estimated network segregation as within-module connectivity, by calculating the module-degree Z score (or within-module strength) ^41,87^ for each brain using the following equation:

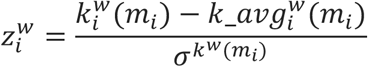

where *m* is the module containing node *i*, 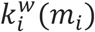 is the within-module degree of *i* (the weighted sum of connections between *i* and all other nodes in *m_i_*), and 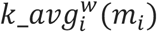 and *σ^k^w^^*^(*m_i_*)^ are the respective mean and standard deviation of the within-module *m*_i_ weighted degree distribution ^41,87^.

### Relating neonatal brain network features with linguistic abilities in childhood

We performed a single holistic, multivariate analysis using PLS regression to simultaneously co-analyze a full set of network nodes metric at the time of normal birth along with sets of later cognitive and linguistic scores. The approach identifies linear combinations of the independent variables that maximally predict linear combinations of the dependent variables ^38,88^.

Prior to the analyses, we first regressed the effect of confound variables on brain and behavior metrics; these were PMA, GA, sex for the network nodes measures; and socioeconomic score (SES), GA, sex for behavioral measures. All predictors and responses were standardized to have zero mean and unit variance. We evaluated different models linking brain network nodes measures with linguistic and cognitive scores. Predictor variables for each model were: (I) streamlines count; (II) weighted degree (ranked; calculated as nodal strength); (III) structural (modal) controllability (ranked); (IV) structural (average) controllability (ranked); (V) network integration (calculated as weighted participation coefficient); (VI) network segregation (calculated as within-module degree z-score); and (VII) atlas-based cortical and sub-cortical volume. All models had the same number of predictor and response variables. Response variables comprised cognitive and linguistic sub-scales of the BSID-III ^58^ (problem solving; receptive language; expressive language) and PARCAR ^59^ (nonverbal cognition; vocabulary; combinatorial language). Linguistic abilities at this age may not be independent of the general cognitive level, and therefore it is key to consider general cognitive development when studying language learning.

The relationship between predictor and response variables was estimated for each model using PLS regression based on the SIMPLS algorithm ^89,90^, with 10-folds cross-validation and 100 Monte-Carlo repetitions. The number of PLS components or latent factors, the linear combinations of the variables in X and Y, was chosen for each model with cross-validation in order to minimize expected error. To estimate the goodness of fit for each model we quantified R^2^, and adjusted-R^2^ (normalizing for sample size and number of components), Mean Squared Error (MSE), Akaike information criterion (AIC), and Bayesian information criterion (BIC), as well as cross-validated predicted R2 and Mean Squared Predicted Error (MSPE). To assess non-parametric statistical significance of each model we performed permutation testing. Response variables were permuted 10,000 times using Permutation Analysis of Linear Models (PALM) ^91^ and each PLS permuted model was fitted with cross-validation and Monte-Carlo repetition as specified above.

For the significant models we characterized PLS components with statistically significant correlation between PLS scores pairs (Family-wise error (FWE) < 0.05, corrected across PLS modes). Where XS are the predictor scores (the PLS components that are linear combinations of the variables in X); and YS the response scores (the linear combinations of the responses with which the PLS components XS have maximum covariance).

For each significant PLS mode, we extracted and normalized the PLS weights (W*), the linear combinations of the original variables that define the PLS component ^38,88^. Given the PLS weight correlation between significant PLS components, we then studied the first principal component of W* across the significant PLS modes in order to derive a summary measure of feature importance for each brain region. The same procedure was repeated to extract the W* from the PLS permuted models in order to calculate the non-parametric statistical significance of each brain region (corrected for FWE across brain regions) ^91^. To aid interpretation only brain regions with significant positive W* were considered (one-tail FWE p-value < 0.05, corrected across regions; corresponding to –log_10_ (p-value) > 1.3010, corrected across regions) ^92^. PLS weights maps not corrected for multiple comparisons are also provided (Fig. S2, S3). PLS coefficients encoding the involvement of behavioral measures in the PLS modes were also extracted and ranked. To aid the interpretation of these coefficients, we also calculated the correlation between each behavioral skill and the first principal component of W* (PLS weights of X) across the significant PLS modes ^93^, thus scaling PLS weights of Y between 0 and 1 (Fig. 2c, 3c).

In order to deliver a more exhaustive characterization of the identified significant model, we also provide further information regarding the relation between predictors (within model) and the PLS model estimation (Fig. S5, S6, S7). All statistical analyses were performed using MATLAB (R2015b, The MathWorks, Inc., Natick, MA, USA).

### Testing the effect of degree of prematurity at birth on brain-behavior links

We removed the effect of all confounds from the dependent and independent variables: PMA and sex from brain metrics; SES and sex from behavior metrics. We then used the same pipeline highlighted before to estimate PLS regression models for both language brain controllability and language brain integration. We tested the effect of premature environmental exposure on the identified brain models of language acquisition by testing whether the PLS scores pairs could discriminate between high vs low GA at birth. We split the sample onto those preterm infants with GA at birth less or equal to 29 weeks (referred as “low” GA group); and those with GA at birth greater then 29 weeks (referred as “high” GA group). A post-hoc data-driven test performed using k-means clustering confirmed the cut-off of 29 GA weeks as the optimal point to divide our sample in two groups. We performed multivariate cross-subject classification based on all estimated PLS scores pairs (or PLS individual subject weights) using cross-validated linear discriminant analysis (LDA) implemented in the FSLNets toolbox for MATLAB. Statistical significance of LDA classification accuracy was calculated with 10,000 permutation of PLS scores pairs.

The effect of postnatal age (or days of ex-utero life) was not considered a variable of interest as it is a linear combination of PMA minus GA and therefore is highly correlated (Fig. S8).

### Testing significance of brain-behavior models against permuted null models and against additional covariates of interest

Aware of the caveats of structural controllability and network neuroscience ^33^, we also assessed the statistical significance of our brain-language results using random null models. For 1,000 times, for each participant, we randomized the individual structural brain network while simultaneously preserving the degree-, weight- and strength-distributions using the function *null_model_und_sign* from the Brain Connectivity Toolbox ^41^. We then fed these matrices to the same pipeline and quantified PLS model goodness of fitting for random networks. This allowed us to compare the “genuine” models of language brain network controllability and integration against the respective random null models’ distributions and calculate the non-parametric random null models-based statistical significance.

## Acknowledgements

We are grateful to the families, clinicians and investigators who made the ePrime study possible, particularly Denis Azzopardi, Mary Rutherford and Maggie Redshaw. We also thank Antonis Makropoulos. The work summarizes independent research supported by the National Institute for Health Research (NIHR) under its Programme Grants for Applied Research Programme (Grant Reference Number RP-PG-0707-10154). It was supported by the Medical Research Council (UK) (MR/K006355/1 and MR/L011530/1), a MRC Clinician Scientist Fellowship to TA (MR/P008712/1) and PhD studentship to PS. The views expressed are those of the authors and not necessarily those of the NHS, the NIHR or the Department of Health.

## Author contributions

P.S., S.J.C., A.D.E. and T.A. conceptualized the study. P.S., J.D.T., T.A. and S.J.C. contributed to image processing. S.F. and A.C. carried out the behavioral data acquisition. P.S., and D.V. designed and carried out the statistical analyses. P.S. wrote the manuscript, with contributions from all other authors. S.J.C., A.D.E. and T.A. have jointly supervised the work.

## Supplementary information

### Supplementary results

#### Alternative models of language brain acquisition

We investigated distinct brain network metrics as alternative mechanisms for explaining language brain acquisition. These alternative models were assessed with the same pipeline as above. We tested the following atlas-based brain metrics: streamlines count; weighted degree (ranked; calculated as nodal strength); structural (average) controllability (ranked); network segregation (calculated as within-module degree z-score); atlas-based cortical and sub-cortical volume. We found that none of these alternative models significantly linked neonatal brain architecture and language abilities in childhood (Table S2). Furthermore, the two models on interest (language brain (modal) controllability and network integration), remained significant even after Bonferroni correction across models.

#### Quantification of PLS model significance against random null models

Aware of the limitations and caveats of structural network controllability ^33^, and more in general of graph theoretical measures, we also tested the identified significant models against non-parametric permutation-based null network models. For 1,000 times, for each participant, we randomized the individual structural brain network while simultaneously preserving the degree-, weight- and strength-distributions. We then fed these matrices to the same pipeline as above and quantified PLS model goodness of fitting for random networks. This allowed us to compare the “genuine” models of language brain network controllability and integration against the respective random null models’ distributions. Crucially both models resulted to significantly relate brain-behavior pairs even against random null models (Table S3).

#### Testing the effect of additional covariates of interest in the identified brain network models of language acquisition

We performed additional tests to understand whether in-scanner head-motion and cortical brain volume could have driven the estimation of brain network models of language acquisition. After regressing these variables into network connectivity, we show that models’ cross-validation remained significant thus demonstrating that neither in-scanner head motion or cortical and sub-cortical volume was driving the estimation of brain network models of language acquisition (Table S4).

## Supplementary figure captions

**Figure S1:**
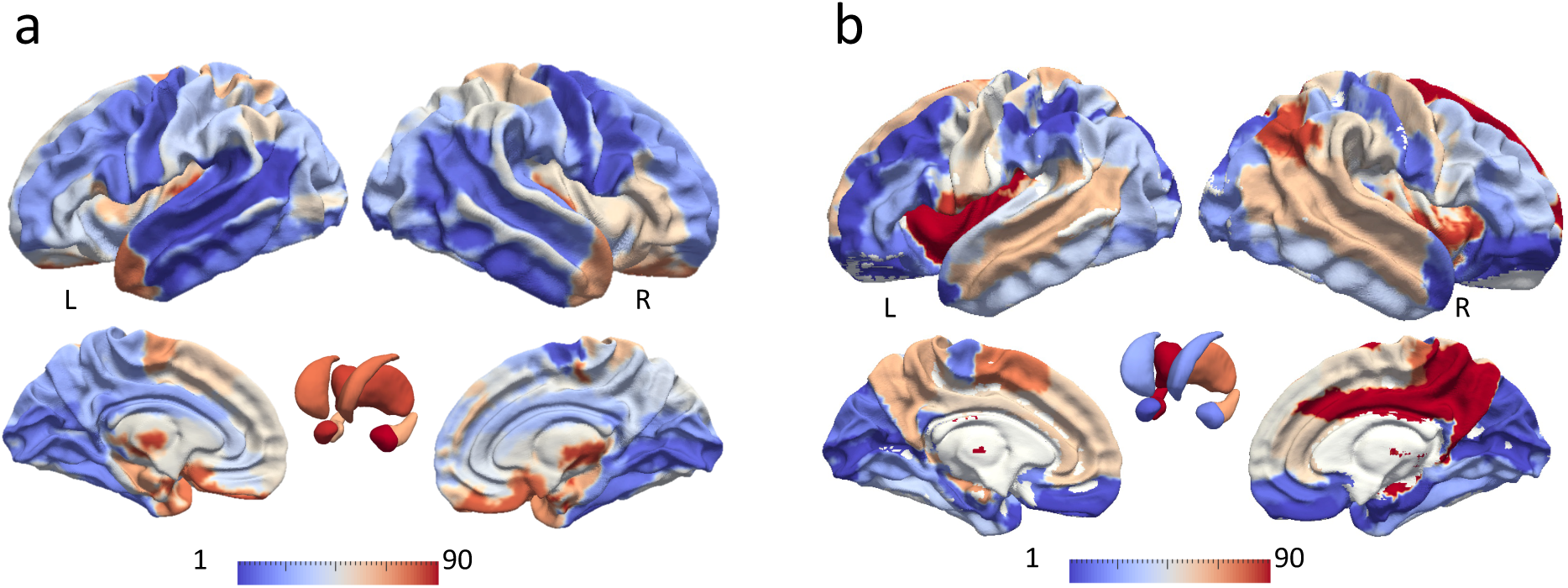
Structural network controllability and integration in the neonatal human brain. We studied the computational properties of the neonatal human brain network. Here, respectively for structural network controllability **(a)** and network integration **(b)**, we are showing normalized group-average ranked values plotted on a surface visualization. Structural network (modal) controllability theoretically predict the ability of a brain region to guide the system’s dynamics. Network integration estimate the ability to rapidly combine specialized information from distributed brain regions.

**Figure S2:**
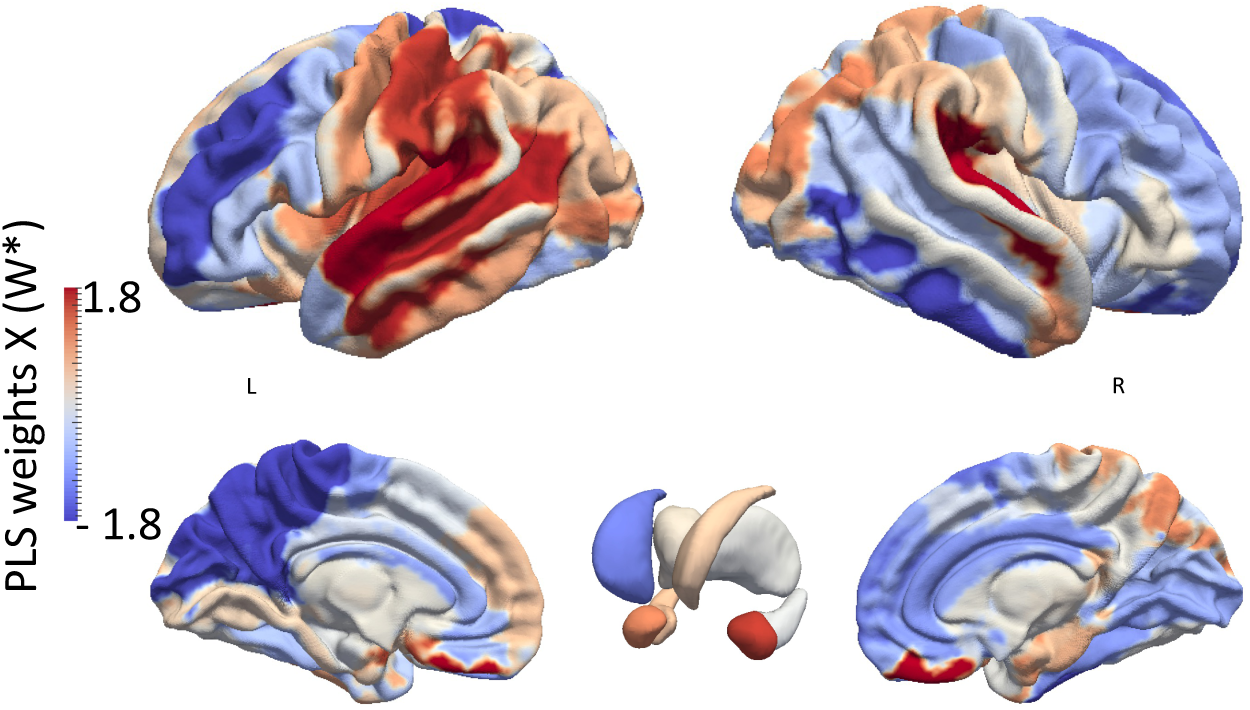
Non-corrected PLS weights maps from language brain controllability. For illustration purposes, we also provide a surface rendered map of the PLS weights (W*) not corrected for multiple comparisons. For multiple-comparisons corrected PLS W* map please refer to Figure 2a.

**Figure S3:**
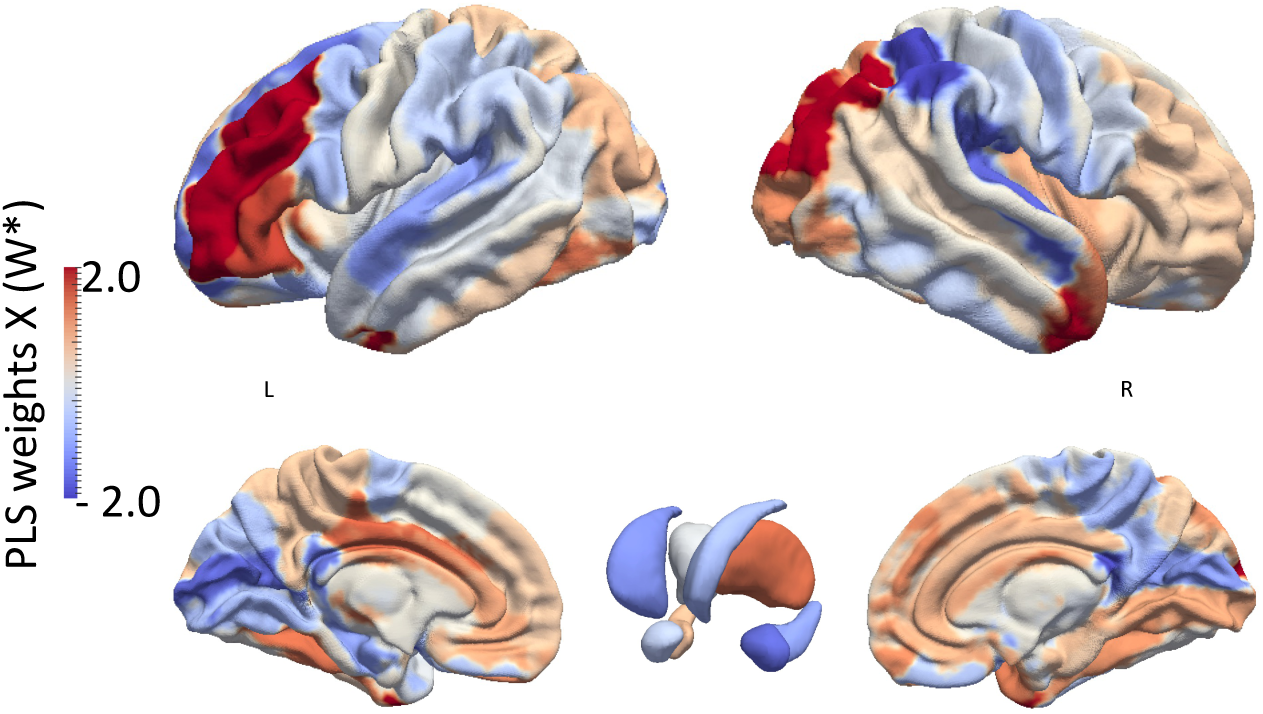
Non-corrected PLS weights maps from language brain integration. We provide a surface rendered map of the PLS W* not corrected for multiple comparisons for the model of language brain integration. For multiple-comparisons corrected PLS W* map please refer to Figure 3a.

**Figure S4:**
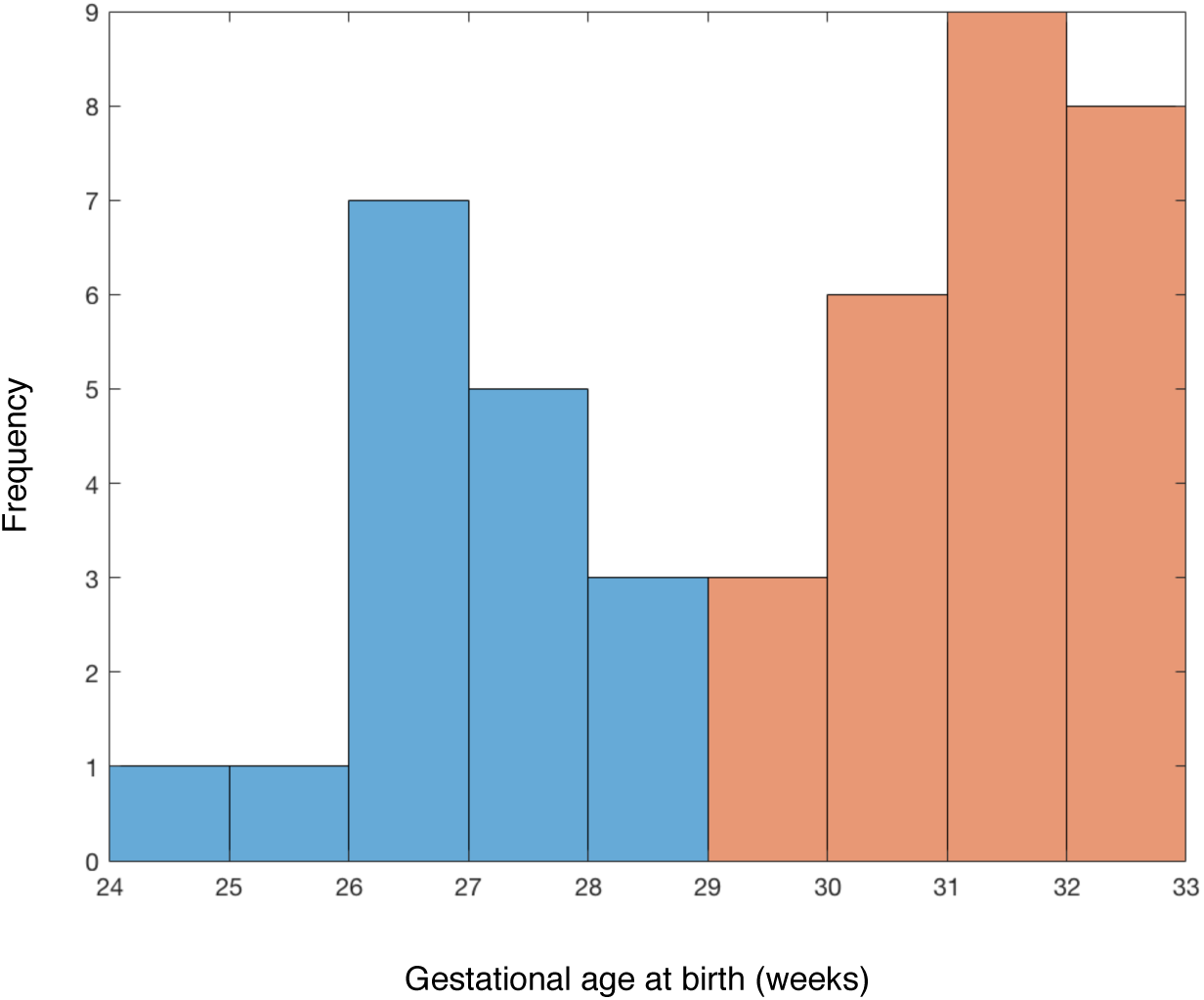
Histogram of gestational age at birth. In blue: sample with low GA at birth (lower or equal to 29 weeks). In red: sample with high GA at birth (higher than 29 weeks). This cut-off choice was confirmed by a post-hoc, data-driven k-means clustering analyses as the optimal value to divide the sample into two groups.

**Figure S5:**
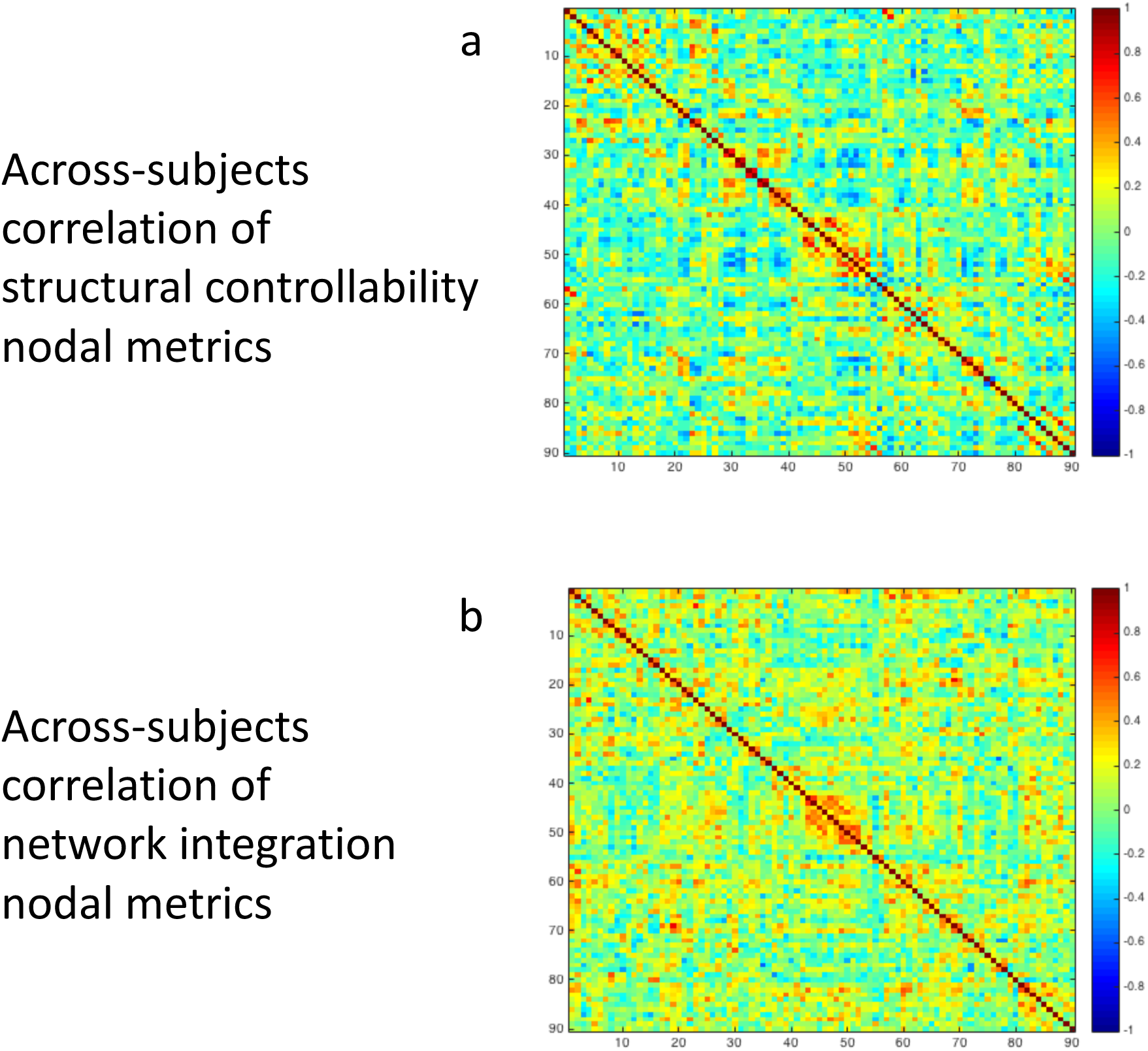
Between-subjects correlation of network metrics are diagonally dominant. Separately for language brain controllability **(a)** and language brain integration **(b)**, here we show the cross-correlation matrices of the network metrics used as PLS predictors. Notably these are diagonally dominant: each feature is more identical to itself when compared to other features. Thus indicating that these sets of predictors carry great variance.

**Figure S6:**
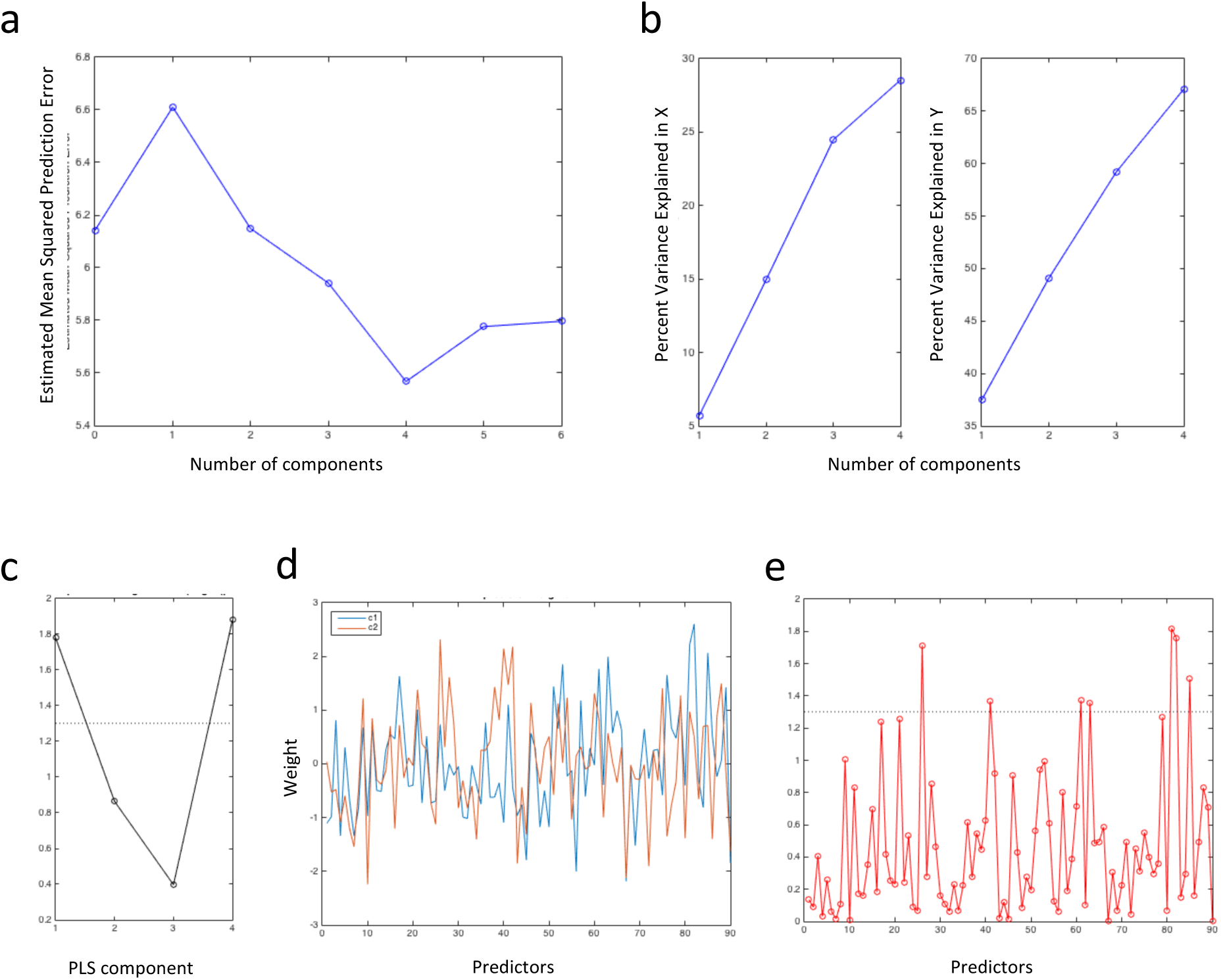
Further results from PLS model estimation for language brain controllability. **(a)** PLS mode number selection based on cross-validated Mean Squared Prediction Error; **(b)** percent of explained variance for the original predictors and responses; “Manhattan” plot of PLS components corrected for FWE across components (statistical significance was set at 1.3010 corresponding to −log10 (0.05)); **(d)** PLS X weights (W*) for each significant PLS component. (Each color represents the PLS W* for a specific PLS component). Given the correlation of W* between significant PLS components, W* were combined across PLS components through one principal component in order to extract summary PLS W*. **(e)** “Manhattan” plot of summary PLS W* corrected for FWE across brain areas.

**Figure S7:**
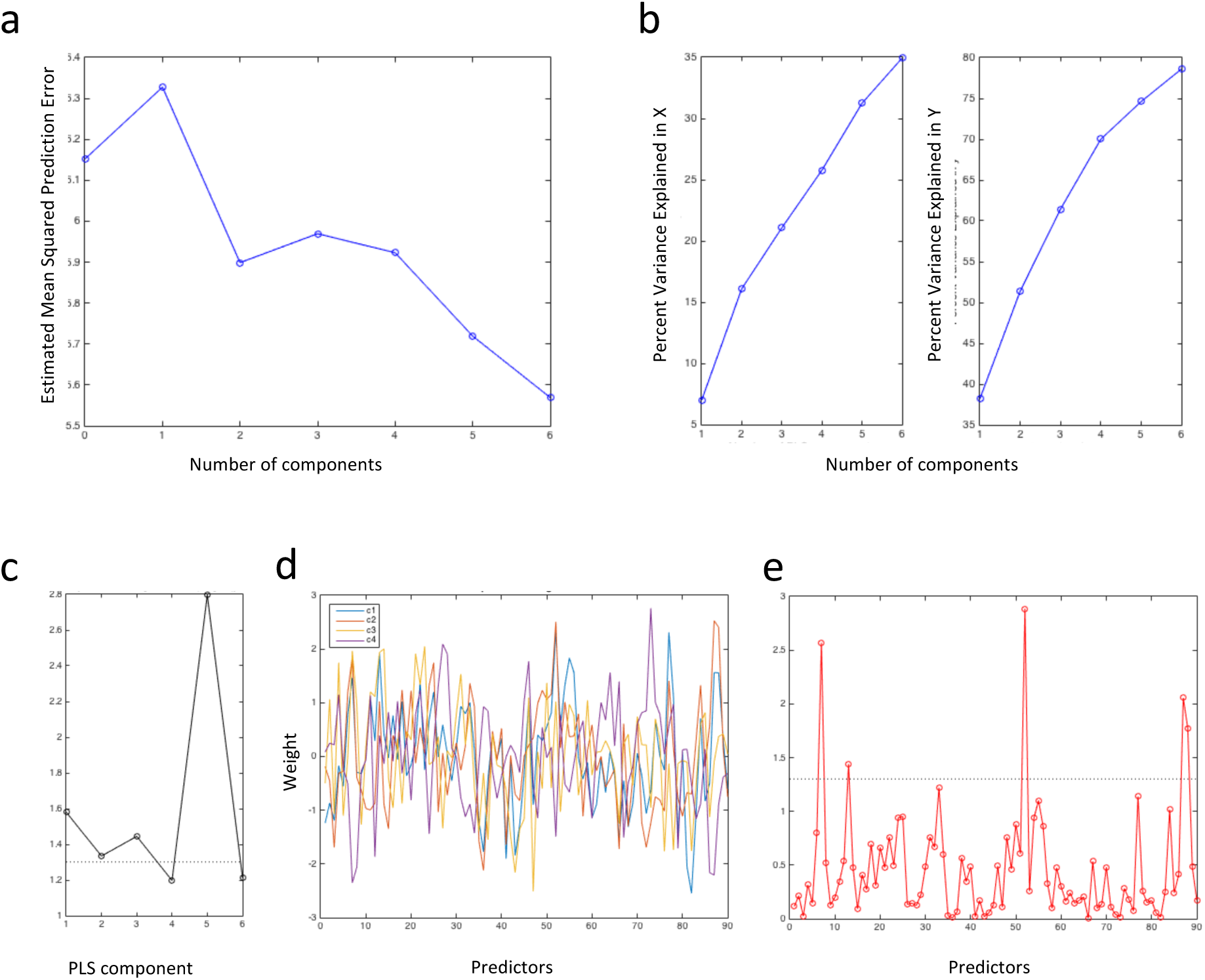
Further results from PLS model estimation for language brain integration. **(a)** PLS mode number selection based on cross-validated Mean Squared Prediction Error; **(b)** percent of explained variance for the original predictors and responses; **(c)** “Manhattan” plot of PLS components corrected for FWE across components (statistical significance was set at 1.3010 corresponding to −log10 (0.05)); **(d)** PLS X weights (W*) for each significant PLS component. (Each color represents the PLS W* for a specific PLS component). **(e)** “Manhattan” plot of summary PLS W* corrected for FWE.

**Figure S8:**
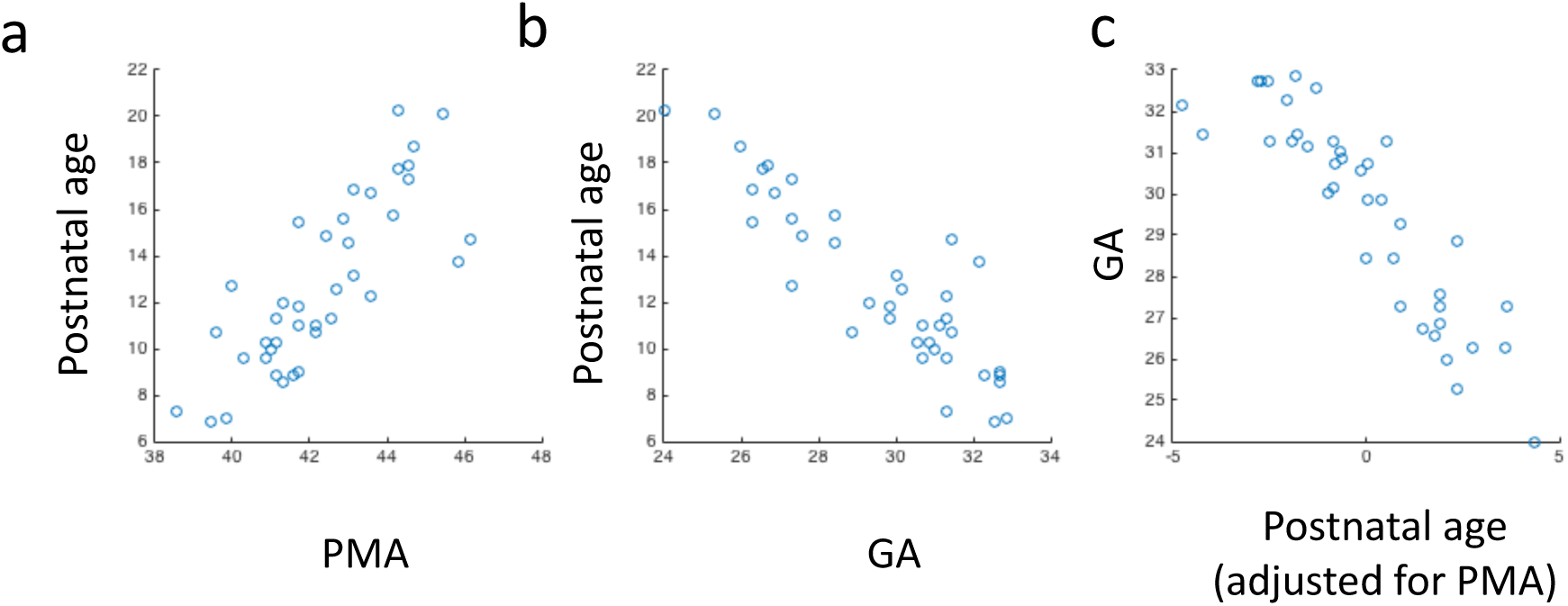
Scatter plots of the relation between postmenstrual age at scan, gestational age at birth, and postnatal age. Postnatal age (or days of ex-utero life) was not considered a variable of interest as it is a linear combination of postmenstrual age minus gestational age and therefore highly correlated.

## Supplementary table captions

**Table S1:**
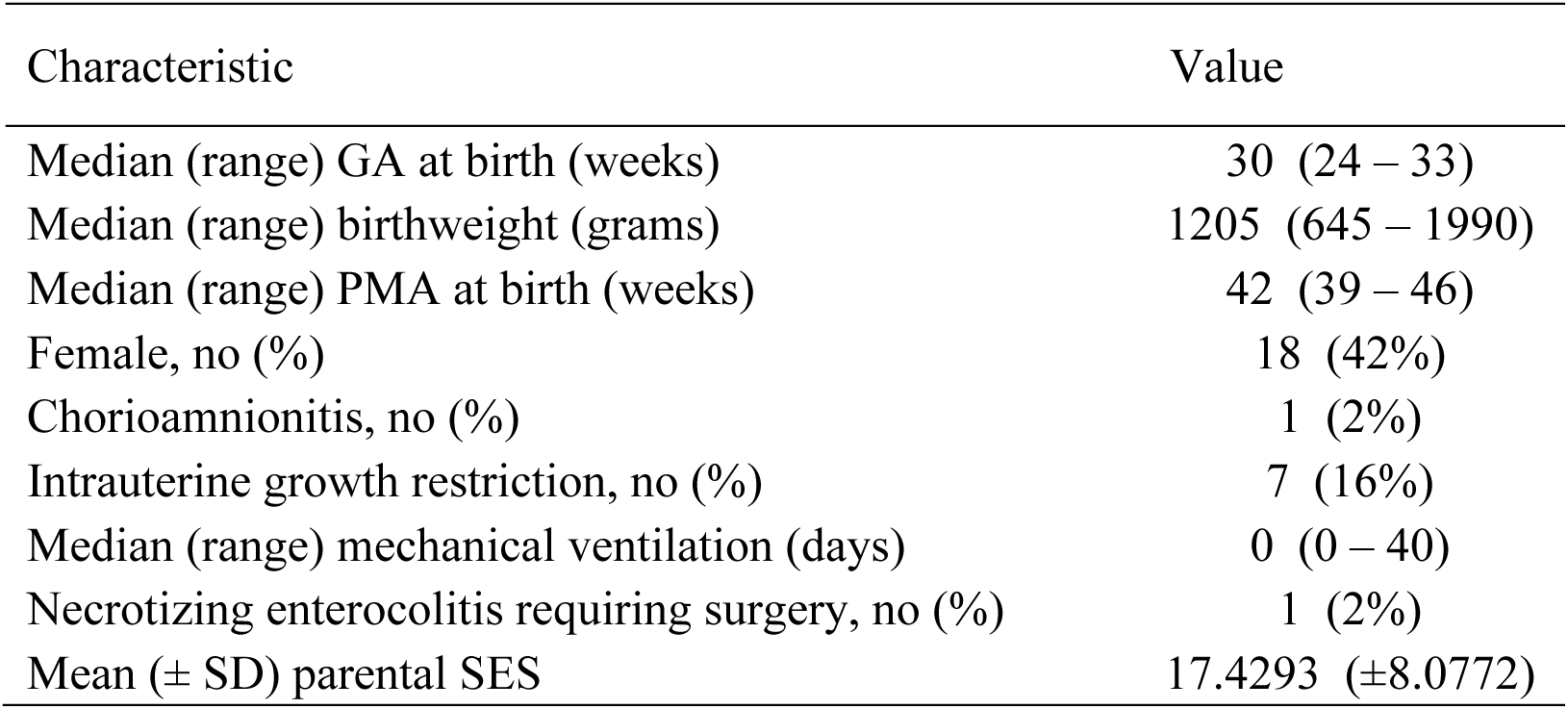
Infants characteristics. This group of preterm born infants has been already object of investigation in a previously published article ^8^.

**Table S2:**
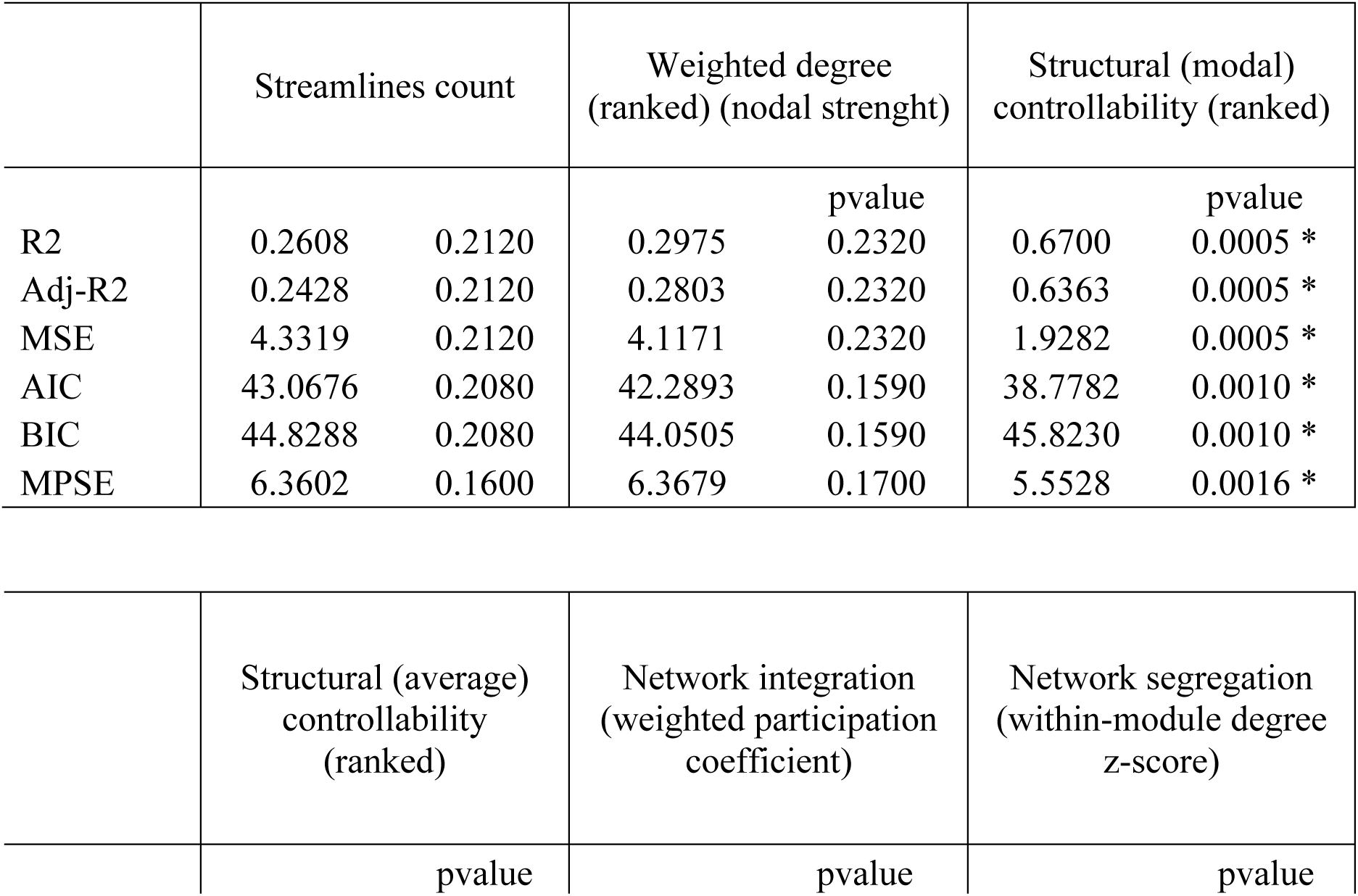
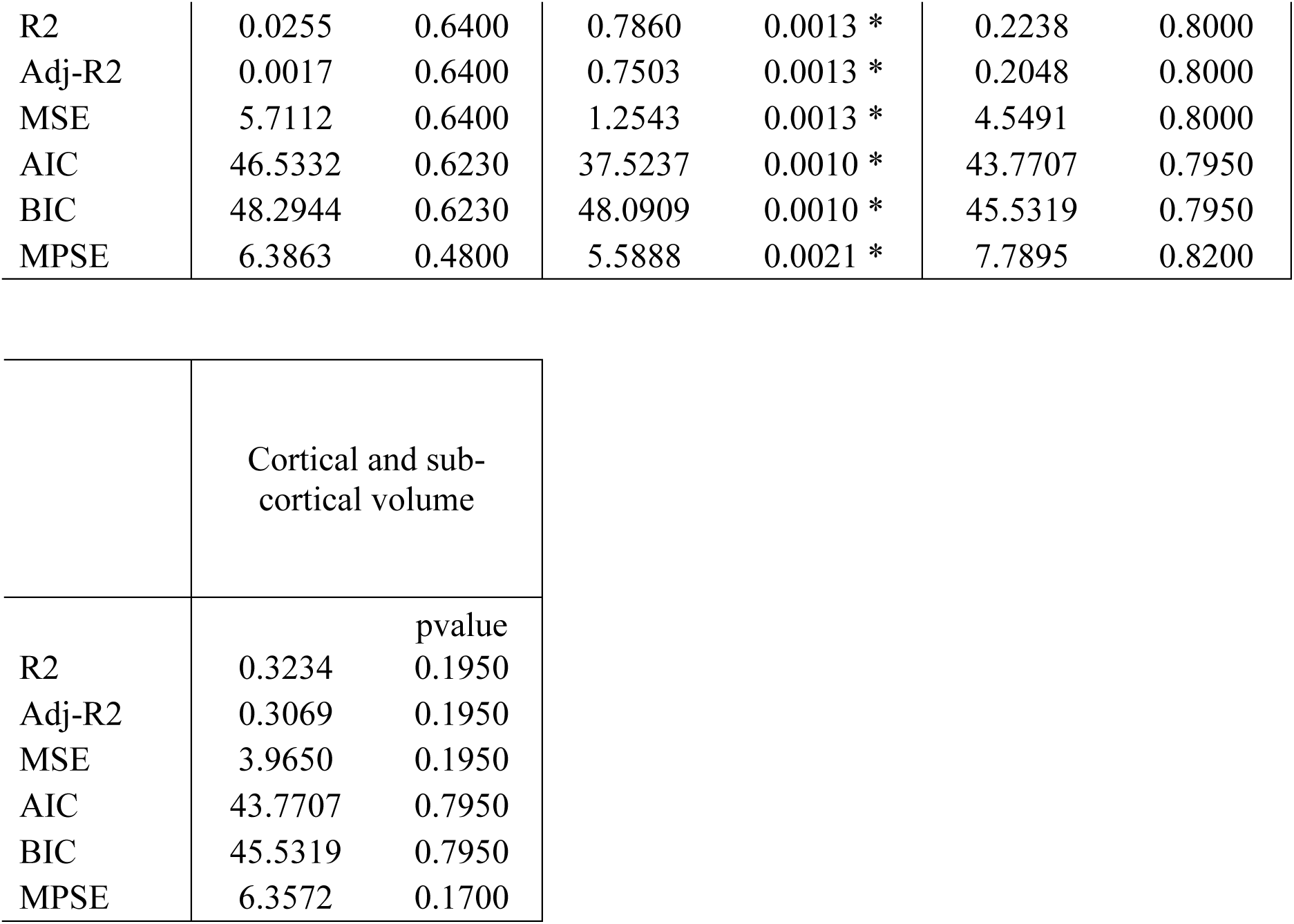
Comparison between alternative models of language brain acquisition. We report Adj-R^2^, MSE and MPSE values for all models tested. All models were tested using PLS regression. They were all based on the same cross-validated pipeline, with Monte-Carlo repetition, and response variable permutations; based on the same set of response variables and on an equal number of predictors. Only the models of language brain controllability and integration resulted to be significant (*) (even after Bonferroni correction across models) and had lowest MPSE and Akaike information criterion values.

**Table S3:** Statistical significance against random null networks. Significant models of language brain controllability and integration were also compared against random null models. For 1,000 times, for each participant, the individual structural brain network was randomized while simultaneously preserving the degree-, weight- and strength-distributions. These individual random null networks were then linked to linguistic and cognitive measures in childhood using PLS regression. Genuine language brain controllability and integration were then compared against the estimated fitting distributions of random null networks in order to calculate statistical significance.

|  |  |  |  |  |
| --- | --- | --- | --- | --- |
|  | Structural (modal)<br>controllability against<br>random null models |  | Network integrations<br>against random null<br>models |  |
|  |  | pvalue |  | pvalue |
| R2 | 0.6700 | 0.0010 | 0.7860 | 0.0010 |
| Adj-R2 | 0.6363 | 0.0010 | 0.7503 | 0.0010 |
| MSE | 1.9282 | 0.0010 | 1.2543 | 0.0010 |

**Table S4:** Model testing against additional covariates of interest. Brain network features of structural controllability and integration were additionally regressed for *in-scanner head-motion* **(a)** and *cortical brain volume* **(b)**. Using the same cross-validated model estimation pipeline we show that prediction results remained statistically significant, thus further demonstrating the robustness of the identified models from confounds of interest.

|  | Structural (modal)<br>controllability (ranked) |  | Network integration<br>(weighted participation<br>coefficient) |  |
| --- | --- | --- | --- | --- |
|  |  | pvalue |  | pvalue |
| R2 | 0.6866 | 0.0010 | 0.7861 | 0.0010 |
| Adj-R2 | 0.6536 | 0.0010 | 0.7504 | 0.0010 |
| MSE | 1.8367 | 0.0010 | 1.2538 | 0.0010 |
| AIC | 38.3759 | 0.0010 | 37.3942 | 0.0010 |
| BIC | 45.4207 | 0.0010 | 47.9614 | 0.0010 |
| pred-R2 | 0.2868 | 0.0020 | 0.2772 | 0.0030 |
| MPSE | 5.4730 | 0.0020 | 5.5096 | 0.0030 |

(b)
|  | Structural (modal)<br>controllability (ranked) |  | Network integration<br>(weighted participation<br>coefficient) |  |
| --- | --- | --- | --- | --- |
|  |  | pvalue |  | pvalue |
| R2 | 0.6410 | 0.0070 | 0.7527 | 0.0320 |
| Adj-R2 | 0.6033 | 0.0070 | 0.7114 | 0.0320 |
| MSE | 2.1036 | 0.0070 | 1.4496 | 0.0320 |
| AIC | 39.8814 | 0.0040 | 39.2055 | 0.0170 |
| BIC | 46.9262 | 0.0040 | 49.7727 | 0.0170 |
| pred-R2 | 0.2427 | 0.0030 | 0.2066 | 0.0030 |
| MPSE | 5.6397 | 0.0030 | 5.7725 | 0.0030 |

